# Modelling surface color discrimination under different lighting environments using image chromatic statistics and convolutional neural networks

**DOI:** 10.1101/2022.11.02.514864

**Authors:** Samuel Ponting, Takuma Morimoto, Hannah Smithson

## Abstract

We modeled discrimination thresholds for object colors under different lighting environments [1]. Firstly we built models based on chromatic statistics, testing 60 models in total. Secondly we trained convolutional neural networks (CNNs), using 160,280 images labeled either by the ground-truth or by human responses. No single chromatic statistics model was sufficient to describe human discrimination thresholds across conditions, while human-response-trained CNNs nearly perfectly predicted human thresholds. Guided by region-of-interest analysis of the network, we modified the chromatic statistics models to use only the lower regions of the objects, which substantially improved performance.

## 1. Introduction

Humans can discriminate very subtle color differences. Mechanisms underpinning color discrimination have been a major focus of past color vision studies, and the non-uniformity of chromatic sensitivities across color spaces has been painstakingly specified in remarkable detail (e.g., [2-5]). These efforts not only informed psychophysical research but have also been used for applications in the display, printing, or textiles industries where knowledge about the discriminability of a given color pair is highly beneficial. However, these early works on color discrimination used colored lights or spatially-uniform color patches as experimental stimuli, and did not directly address the problem of object colour perception. For pigmented objects, spatial variation in pigmentation across the object’s surface introduces spatial colour variation, and objects cannot therefore be characterized by a single color [6]. Acknowledging this limitation, there have been recent attempts to investigate how well our visual system discriminates among natural objects by their color. Hansen et al. [7] directly measured discrimination thresholds using natural objects such as fruits and vegetables and chromatic textures that exhibit spatial color variation. It was found that discrimination ellipses were highly elongated along the axes of maximum variation in the chromatic distributions of the stimuli. Giesel et al. [8] additionally measured thresholds for chromatically variegated stimuli, and they found that discrimination models with eight parameters (four cardinal directions and four intermediate directions) better account for the obtained threshold patterns than four parameter models (cardinal mechanisms alone). In addition to these studies on color discrimination a few studies investigated the color appearance of chromatically variegated objects. One of these color matching experiments showed that humans match to different colors depending on the region of the objects that was specified to be judged [9]. Humans can effectively take the spatial average of hue [10], and mean hue value may indeed determine the overall impression of color for inhomogeneous objects [11].

These works emphasized the importance of specifically considering object-based color discrimination. Moreover, since most objects are visible only by virtue of the light they reflect, and diffuse and specular surface reflections have different imaging geometries, there is additional spatial variation across the proximal image of an illuminated object that derives from the lighting environment and viewing geometry. Natural environments introduce additional complexity because the proximal images of identical objects can exhibit very different color distributions when placed under different lighting environments. Moreover, even within a single object, different regions of the object will receive different illumination, and it is a curious empirical question whether we might correct them differently [12]. Yet the influence of lighting environments on color discrimination has been little studied, especially in the presence of complex object properties (e.g., three-dimensionality and specular reflection). To tackle this, Morimoto & Smithson [1] measured chromatic thresholds using glossy and matte objects, placed under one of three lighting environments. Results showed that discrimination ability is strongly influenced by the chromatic distributions of the lighting environments in which objects are placed (Figure 1). Also, discrimination thresholds were generally higher for glossy objects than matte objects. However, the 2018 paper did not directly address what kind of signals in the proximal images of the objects lead to these different discrimination performances.

**Fig. 1.**
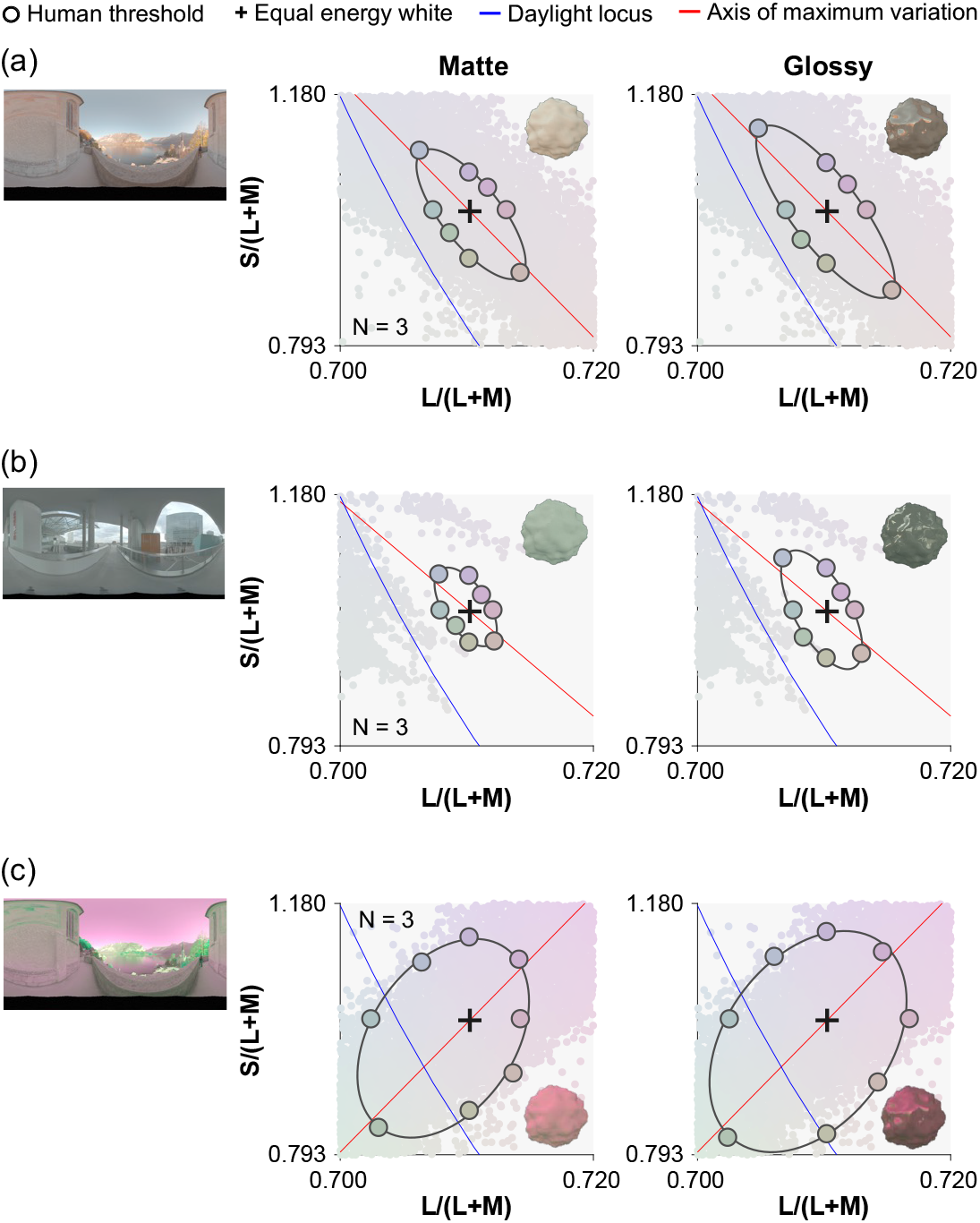
Human discrimination thresholds collected in a previous study [1] for (a) environment 1, (b) environment 2, and (c) environment 3. Left pictures show environmental illumination [13], and the two plots show discrimination thresholds for matte (left) and glossy (right) objects, averaged across three participants. The small colored dots in the background of each plot show the color distributions of each lighting environment. Environment 3 was generated by inverting the chromatic distribution of environment 1 along the S/(L+M) axis.

Thus, the aim of this study was to uncover potential strategies participants took in performing surface color discrimination of complex objects. We tackled this using two distinct computational modeling approaches. For the first set of models, each model uses one of 60 pre-defined chromatic statistics directly computable from the proximal image of a given object and makes discriminations by comparing these statistics across objects. In a second approach, instead of specifying what the model should do, we trained convolutional neural networks (CNNs) on chromatic discrimination tasks using 160,280 images. Importantly, for each CNN images were labeled either by their physical ground-truth labels or by perceptual labels. We formally evaluated the performance of each model (i.e. those based on chromatic statistics, and each of the trained networks) using the adaptive staircase procedure that tested human participants in the previous study. Finally, we used occlusion sensitivity maps to reveal the potential strategies used by the CNNs to solve the discrimination task. Then, guided by this suggested strategy, we developed models that segmented the images before computing specified chromatic statistics.

## 2. Method

### 2.1 Modeled datasets and experimental task from the previous study

The psychophysical data modeled in this paper were all collected in a previous study [1]. Details are provided in the previous study, but here we briefly introduce the experimental design and stimuli. Thresholds were measured along eight hue directions from the equal energy white point, using an adaptive staircase procedure [14]. Test images were generated by rendering bumpy spheroid stimuli placed under one of three lighting environments (shown in Figure 1, obtained from a publicly available database, http://dativ.at/lightprobes/). The first two lighting environments featured chromatic distributions that broadly corresponded with the daylight locus [15], while the third environment was created by chromatically inverting the first. Surface roughness of the object was fixed at 0.2, and the object was either matte or glossy (specularity 0.20 in the Ward reflectance model [16]). All experimental stimuli were generated by a physically based renderer Mitsuba [17] and RenderToolBox3 [18]. We performed multispectral rendering from 400 nm to 700 nm with a 10 nm step size. The resultant spectral images were converted to LMS images (corresponding to the signals in the long-, middle- and short-wave sensitive human cones) using Stockman & Sharpe’s cone fundamentals [19].

There were six experimental conditions (two levels of glossiness × three lighting environments). For each trial, we simultaneously presented four objects rendered under the same lighting environment in a square configuration. The three distractor objects were assigned an achromatic surface colour (flat reflectance), while the fourth stimulus – the target – was assigned a reflectance biased in one of the eight directions of hue angle. Participants were asked to select the odd-one-out object in terms of surface color. In every trial the camera angle for each object was randomly assigned from six possible angles (0° to 300° in 60° steps), and we also ensured that within each trial, no two stimuli shared the same camera angle.

### 2.2 Computational modeling

#### 2.2.1 Chromatic statistics models

As shown in Figure 2 (a), we constructed model observers that base their judgment on specified chromatic statistics. We selected 60 candidate statistics that are directly computable either in the MacLeod-Boynton chromaticity diagram [20] or in *L*\**a*\**b*\* color space, which are physiology-based or perception-based spaces, respectively. Chromatic coordinates *L*/(*L*+*M*) and *S*/(*L*+*M*) in the MacLeod-Boynton diagram and photopic luminance are collectively termed MB color space in this study.

**Fig. 2.**
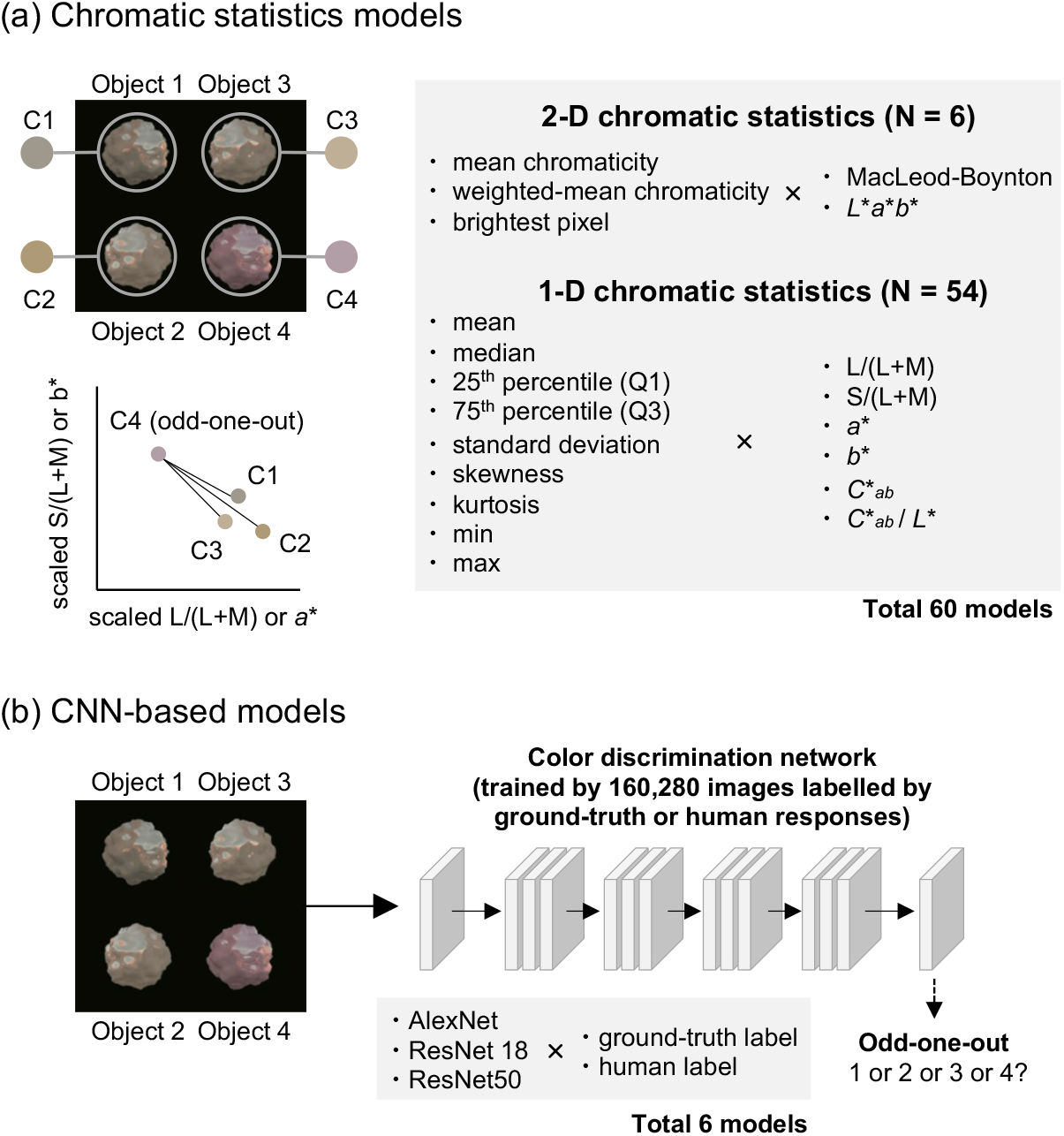
Two modeling approaches used in this study. For (a) chromatic statistics models, we predefined 60 chromatic statistics and for each trial the model extracted the specified statistics from each object (C1 to C4). The model compared the statistics and identified the odd-one-out. In contrast (b) CNN-based models were simply given 160,280 training images labeled either by ground-truth or human response labels. Three network architectures were used, generating six trained CNN models in total.

We used two types of chromatic statistics models. The first models summarize variation of chromaticity over the surface by taking (1) mean chromaticity [21], (2) mean chromaticity weighted by luminance or lightness [22] or (3) the color of the brightest pixel [23] in MB color space or *L*\**a*\**b*\* color space. Thus, a single chromaticity that consists of two scalar values was taken as an estimate of surface color for a given object. These six models are referred to as 2-D models. The second type of model focuses on a single dimension in a color space, namely *L*/(*L*+*M*), (2) *S*/(*L*+*M*), (3) *a*\*, (4) *b*\*, (5) *C*\*_*ab*_, (6) saturation (*C*\*_*ab*_ / *L*\*) and summarizes the spatial variations of these by taking (1) mean (2) median, (3) 25th percentile value, (4) 75th percentile value, (5) standard deviation, (6) skewness, (7) kurtosis, (8) minimum value or (9) maximum value. The combination of these (six dimensions × nine statistics) resulted in 54 models which were termed 1-D models in this study. Some of these 1-D statistics have been used as candidate statistics to estimate surface color from natural objects [11]. To convert XYZ to *L*\**a*\**b*\*, we defined a white point separately for each lighting environment. We first rendered 12 white matte objects whose diffuse reflectance are 100% across visible wavelengths from 0° to 330° in 30° steps, and then we computed average XYZ values over the object and over 12 camera angles. By defining the white point per environment in this way, the *L*\**a*\**b*\* values (and statistics based on them) largely compensate for differences in the global means of the chromatic statistics in the proximal images of the objects. We also note that it is unlikely that participants actually make judgments based only on, for example, *a*\* values, but these simulations are intended to test the extent to which such reduced models can perform the discrimination task.

In each trial a given model calculated the specified statistic for each of the four stimuli and picked the odd-one-out by selecting the object that has the maximally different chromatic statistic from the other three objects, assessed by a sum of three differences. For the models based on 2-D statistics, we used the 2-D distance on the chromatic plane of MB space or *L*\**a*\**b*\* space (see lower left plot in Figure 2 (a)). When calculating the distance in MB space, the two axes were scaled by the discrimination thresholds (previously measured using homogenous colored discs) to roughly equate their scales. Threshold estimates from each model were derived using an adaptive staircase to test the modeled ‘observer’ as human observers had previously been tested psychophysically (see below).

Previous studies that used computational observer models typically introduced noise in the estimation of the color of the surface or illuminant (e.g. [24]), but in this study no noise was added to estimation of chromatic statistics for two reasons. First, we generally found that thresholds obtained from these models were larger than thresholds found from humans. Thus, adding noise did not improve the model’s ability to predict human thresholds. Second, we used chromatic statistics derived from different color spaces, and it was difficult to find the equivalent strength of noise across the color spaces, which would likely confound the effectiveness of each chromatic statistic.

#### 2.2.2 CNN-based models

The second set of models we tested were convolutional neural networks (CNNs) as shown in Figure 2 (b). We selected three widely used CNN architectures for object recognition tasks: AlexNet [25], ResNet18, and ResNet50 [26]. Each model was trained in two ways, either with the ground-truth labels (to optimize objective performance of the models), or with human perceptual labels (to train the model to reproduce human-like chromatic thresholds). The second approach relied on data from the previous study in which a large number of images were labeled by the object location that human participants selected during each trial of the experiment. The previous study used an adaptive staircase procedure, requiring 10 to 40 trials for each threshold estimate, with stimulus magnitude typically clustered around the threshold. Putting all repetitions, conditions and participants together, the total number of images presented during the experiment summed to 20,070 images. To increase the training set we applied data augmentation by a factor of eight, based on two assumptions: participants give the identical response even when (1) a given image is horizontally mirrored, and (2) the target object was presented at any of the other three positions. The resulting total of 160,280 images were labeled either by human responses (the location selected by human participants in the previous study) or by physical ground-truth (correct location of target object from 1 to 4). Each pixel stored LMS cone signals instead of RGB values to make sure that networks will receive the same sensory inputs as humans. Each network was trained from scratch using a stochastic gradient descent with momentum algorithm with a fixed learning rate of 0.0003 for 10 epochs by which point performance of the network plateaued. The batch size was 64, and the Deep Learning Toolbox in MATLAB was used.

### 2.3 Estimation of chromatic thresholds from models

We formally estimated chromatic thresholds from all models using new image datasets in which all stimuli were rendered from different camera angles (30° to 330°, every 60° steps) from the training dataset. Thresholds were estimated using a Palamedes adaptive staircase [14], starting from large differences in saturation between the target and distractor stimuli, and decreasing the saturation difference if the algorithm correctly identified the odd-one-out. The staircase algorithm increased the target stimulus’ saturation if the algorithm identified an incorrect stimulus. Staircases consisted of a minimum of 10 and maximum of 40 trials to improve reliability, and to match the human paradigm. One session consisted of 8 interleaved staircases to measure thresholds for all hue angles. 20 sessions were carried out, such that 20 thresholds were obtained for each data point.

## 3. Results

### 3.1 Model performances

Figure 3 summarizes the root-mean-square-error (RMSE) between model thresholds and human thresholds (averaged across three participants) for all conditions calculated in scaled MB chromaticity space. On the left-hand plots, we show RMSE for all CNN-based models and for the top-10 chromatic statistics models. An RMSE value of 0 denotes a perfect match to human thresholds, with higher values showing a worse match. On the right-hand plots, we show thresholds of the best CNN models (upper-right) and best chromatic statistics (lower-right) by colored symbols and fitted ellipses, along with human thresholds (black symbols and ellipses). These best models are ringed by red circles on the left plot.

**Fig. 3.**
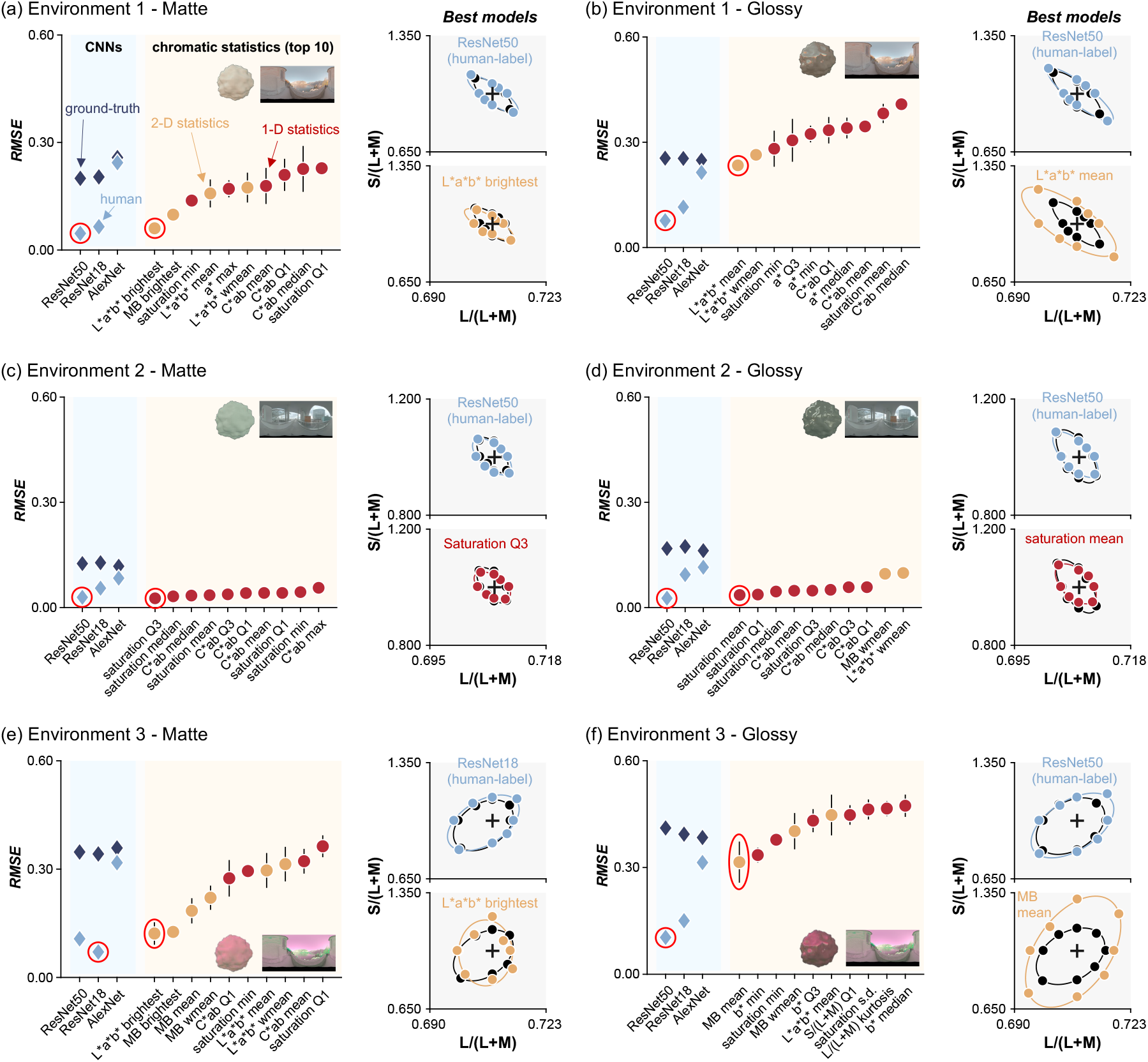
Prediction error (RMSE) between human thresholds and model thresholds for all CNN-based models (light blue and dark blue diamonds in left plot) and for the top-10 chromatic statistics models (orange and red circles) for each condition (a) - (f). Lower RMSE shows better model performance. Estimated discrimination thresholds for the best models (ringed data points) are shown on the right-hand plots.

Looking across panels, it is notable that ResNet50 (or ResNet 18 for (e)) trained by human response labels shows nearly perfect match to human thresholds in all conditions. In contrast, for the chromatic statistics models, the best performance was derived from different models for different conditions.

For environment 1 and matte stimuli (panel (a)), we see that, of the chromatic statistics models, the color of the brightest pixel in *L*\**a*\**b*\* color space best predicts human performance. However, there was a significant difference in performance averaged over 20 sessions between this model and ResNet50 (non-paired two-tailed t-test, *t*(38) = 3.20, *p* = 0.0028). The brightest color in MB color space is ranked second. Both of these summary statistics suggest that humans used color at high intensity points on the image of the object’s surface to perform discrimination for matte stimuli. However, these brightest-point-based models are not listed within the top-10 models for the corresponding glossy condition (panel (b)). Instead, the mean chromaticity or weighted mean chromaticity model in *L*\**a*\**b*\* color space shows the best agreement with human thresholds. Importantly, the thresholds are still substantially larger than human thresholds and ResNet50.

The ranking of chromatic statistics models in environment 2 is very different from environment 1. It is shown that saturation-based methods were ranked first for both matte (panel (c)) and glossy (panel (d)) stimuli. RMSE error for this model is not significantly different from ResNet50 trained on human-labels for the matte condition (non-paired, two-tailed t-test, *t*(38) = 1.54, *p* = 0.132) but is significantly different for the glossy condition (non-paired, two-tailed t-test, *t*(38) = 4.07, *p* = 2.26×10-4).

Data from environment 3 show a similar trend to environment 1 where the brightest color in *L*\**a*\**b*\* color space is the best predictor for matte stimuli (panel (e)), and for glossy stimuli (panel (f)) the mean chromaticity model in MB color space is the best.

CNNs trained on ground-truth labels consistently showed higher RMSE values (with respect to human thresholds) than CNNs trained by human response labels. As shown in Figure 4, this is because these networks can perform the task nearly perfectly, leading to large RMSE values in comparison to the less perfect human thresholds. Since the CNNs were given the correct answer for each trial, it is not surprising that CNNs produce lower discrimination thresholds than CNNs trained by human labels. Importantly, the low thresholds for ground-truth CNNs confirm that the stimuli contain information that allows for near-perfect discrimination, though this information is not available to, or is not correctly utilized by, humans or the chromatic statistics models.

**Fig. 4.**
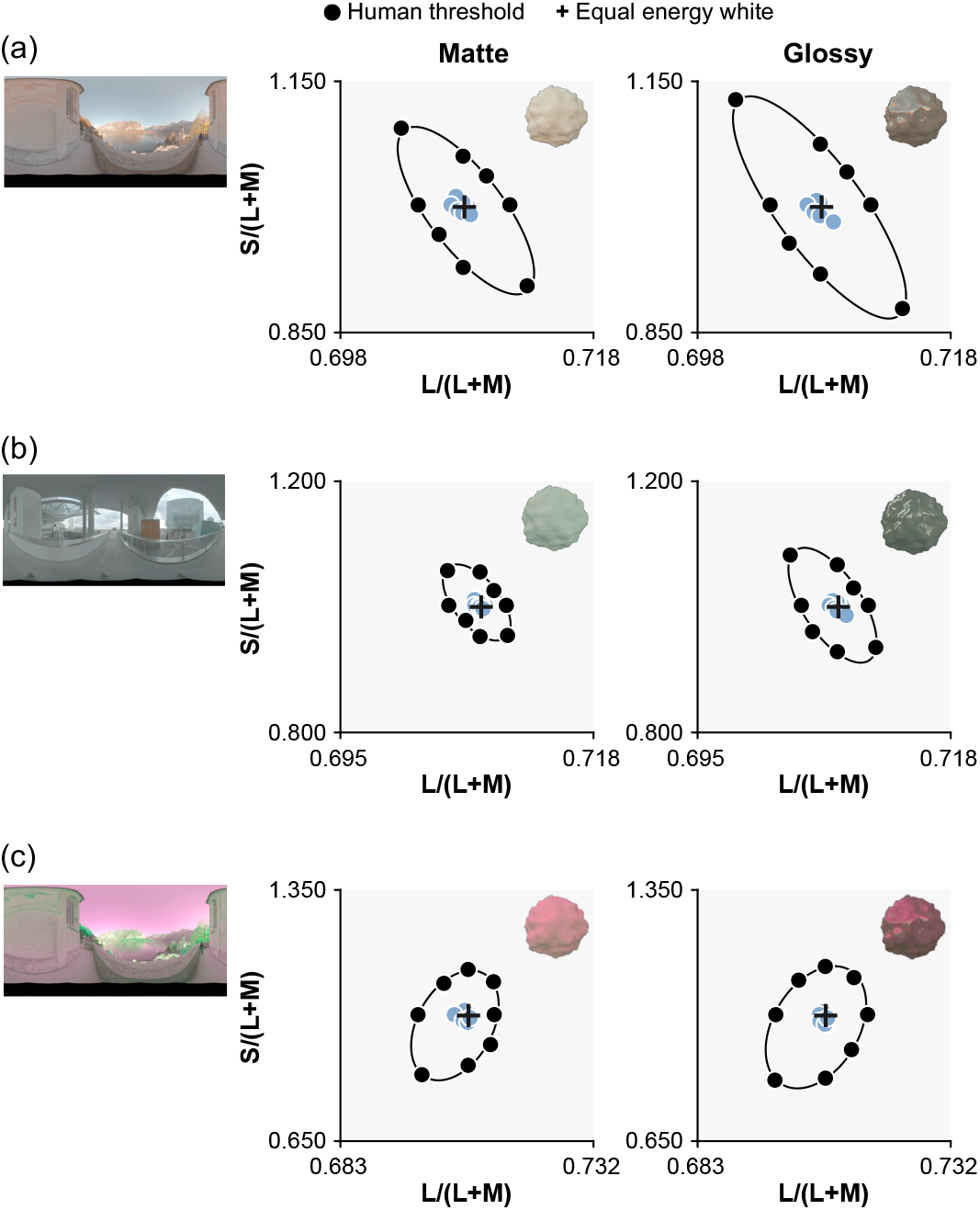
Thresholds derived from ResNet50 trained on ground-truth labels (light blue circles). It is shown that the ground-truth CNN shows much lower thresholds than human psychophysical data (black circles and ellipses).

In summary, these modeling approaches reveal potential strategies that human participants might have used when discriminating surface colors of complex objects within lighting environments. For matte objects presented under environments 1 and 3, human performance is correlated with the model that bases its judgment on the color of the brightest region of the stimuli. For gloss objects, human performance is closer to that predicted from more global statistics such as mean chromaticity or weighted mean chromaticity, since, for these stimuli, the brightest point corresponds to specular reflection, which does not convey information about the surface color of the object.

It is curious that different trends were observed in environment 2, but this could be explained by the relatively small chromatic variation across the proximal image of objects under environment 2. In other words, surface color discrimination in environment 2 was closer to simple saturation discrimination of a uniform surface, which might be why saturation-based metrics were better predictors of the empirical thresholds. To confirm this, we analyzed the standard deviation of L/(L+M) and S/(L+M) values across the proximal images of matte and glossy achromatic objects (same as distractor stimuli) under environment 1 and 2. This calculation was repeated over six different camera angles (from 30 degree to 330 degrees, with 60 steps). For matte objects, the averaged standard deviation of *L*/(*L*+*M*) was significantly lower for the six camera angles in environment 2 than for those in environment 1 (non-paired two-tailed t-test; *t*(10)=14.4, *p* = 5.05×10^−8^), but not for *S*/(*L*+*M*) (non-paired two-tailed t-test; *t*(10)=0.1458, *p* = 0.8870). For glossy objects, the variation in both chromatic axes was significantly lower for environment 2 than for environment 1 (non-paired two-tailed t-test; *t*(10) = 30.9, *p* = 2.95×10^−11^ for *L*/(*L*+*M*); *t*(10) = 5.14, *p* = 4.34×10-4 for *S*/(*L*+*M*)). Thus, the image statistics are broadly consistent with an explanation based on chromatic variation. In addition, it is worth noting that, as seen in Figure 2 of the 2018 paper, the mean chromaticity of environment 2 is biased away from equal energy white towards bluish-green. However, to compute *L*\**a*\**b*\* coordinates, we set the white points for each environment based on the mean chromaticity of objects rendered under that environment, and thus we expect that the chromatic bias of environment 2 was largely discounted in the saturation metric, which thus directly provided information about the saturation of the objects without much interference from the color of the lighting environment.

For the gloss conditions for environment 1 and 3, we found limited applicability of the implemented chromatic statistics models. Nevertheless, ResNet50 trained on human responses provided good fits to the human psychophysical thresholds for these conditions as well. Thus, we thought that if we could understand the potential strategy of this trained CNN to perform the task, it might give us a direct hint as to the strategy adopted by human participants. This notion encouraged us to analyze the regions of interest for this network trained on human responses.

### 3.2 Occlusion sensitivity

We used occlusion sensitivity [27] to reveal regions of the stimuli that were significant to the networks in making their classifications. The basic idea of this technique is to hide a specific region in the image with a square mask and record the change of classification results. This mask is moved across the whole image, and if the area causes a strong change of response, that region can be deemed critical for the discrimination task.

Figure 5 shows occlusion sensitivity maps derived from ResNet50 trained by the human labels and ground-truth labels, for the glossy conditions in each environment. During the analysis, the images included four objects, arranged in a square configuration, but only the target region is shown here. For a given hue angle, stimulus strength (saturation of the surface reflectance) was fixed at twice the ResNet50 thresholds measured in each condition, and separate strength was used for human ResNet50 and ground-truth ResNet50. The results shown here are for hue value 0 degree but we found similar results for other hue angles. It seems that both CNNs – when classifying correctly – were pulling information from specific regions on the lower part of the target stimulus. The region deemed to be useful by the ground truth-trained models was also typically larger than that of the human-trained models, while still being biased towards lower regions of the stimuli. This pattern was also present in ResNet18 which was also relatively successful in predicting human thresholds. Furthermore, we confirmed that matte objects lead to similar maps.

**Fig. 5.**
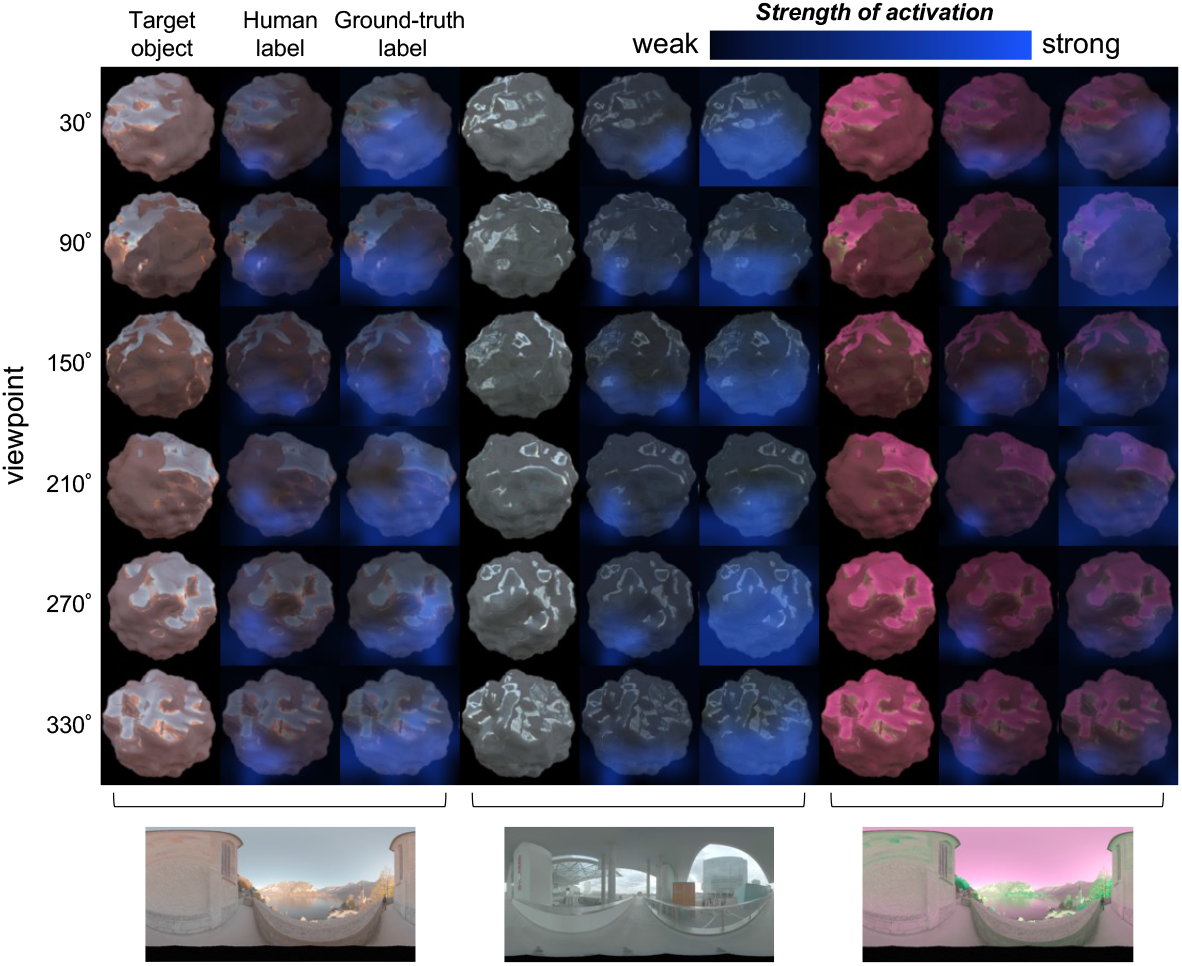
Occlusion sensitivity maps from ResNet50 obtained for objects used in the glossy conditions for different viewpoints under three different lighting environments. For these maps, the hue angle was 0 degrees, and the magnitude of stimuli was fixed at twice the ResNet50 thresholds (separate strength was set for human ResNet50 and ground-truth ResNet50). The input image to the network was four objects located in square configuration (as shown in Figure 2), but only the target region is shown here. For each environment, the leftmost column shows the target object in sRGB format, and the center and rightmost columns show the strength of activation overlaid on the image of the object obtained from ResNet50 trained on human labels and the ground-truth labels, respectively.

To summarize, these analyses suggest that humans might have extracted information from the lower regions of the stimuli to get reliable estimates of surface colour. This observation encouraged us to create a further image-segmentation model which computes chromatic statistics from a restricted region of a given object image.

### 3.3 Image segmentation models

Guided by what we learned from occlusion sensitivity analysis of CNNs, we decided to further implement chromatic statistics models that operate only on pre-determined interesting regions of a given object, which we call image segmentation models. The occlusion sensitivity analysis identified lower regions of the objects. The first set of models simply calculate image statistics from the lower p% height (p= 25, 50, and 75) of the object’s region in the proximal image. An additional characteristic of the lower regions of the objects is that they contain fewer specular highlights. To test whether this was the critical feature of the lower parts of the objects in improving performance, we made a second set of models that take image statistics from pixels that have intensity below p% of the highest intensity pixel over the object’s surface (p = 25, 50 and 75). Since the mean chromaticity and weighted mean chromaticity models were the best predictors of human thresholds for glossy conditions in environment 1 and 3 (Figure 3), we implemented only these chromatic statistics for the image segmentation models.

We found that models that compute statistics only from the lower 25% region of the objects are indeed better predictors of human behavior. Figure 6 is a summary plot showing discrimination thresholds of the best overall models in this study. We found that mean chromaticity or weighted mean chromaticity from the lower 25% of the object region best predicted human thresholds in environments 1 and 3. In supplementary material, we report RMSE values for all models implemented in this study.

**Fig. 6.**
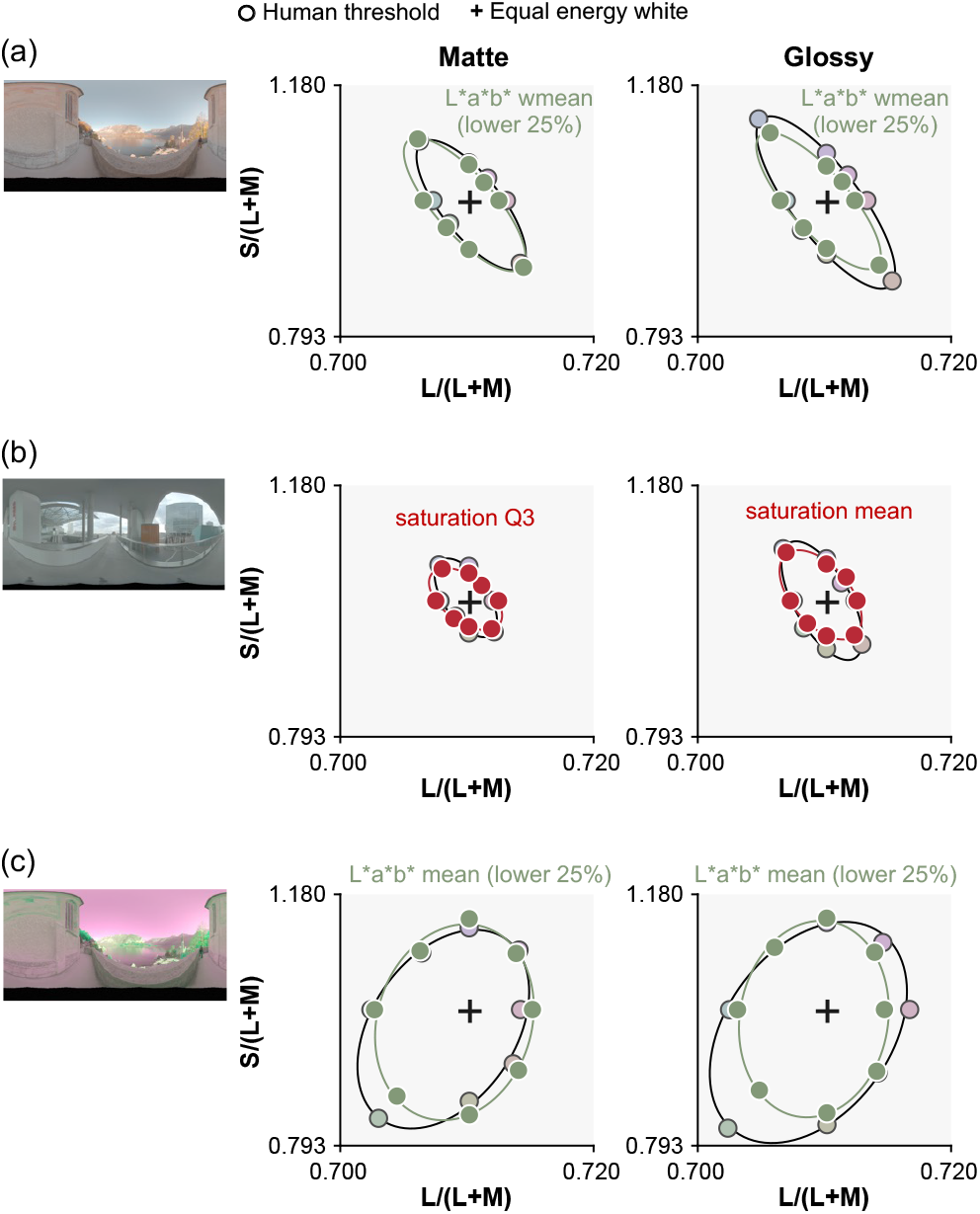
Overall best models for each condition. Using chromatic statistics computed from lower part of the object yielded the best prediction in (a) environment 1 and (c) environment 3 while a saturation metric (C*ab / L*) yielded the best prediction in (b) environment 2, in which chromatic variation around the white point introduced by the lighting environment was smaller than in the other two environments.

It is interesting that a model that computes statistics from the lower part of the object showed more similar thresholds to those of human observers than a model that uses only dim pixels. Computing the statistics on the dim pixels would remove specular highlights. Similarly, there are fewer specular highlights in the lower parts of the object images. But, computing statistics from lower regions of the object is different from using dim pixels since the lower part is shadowed, meaning that it does not receive light from the brighter regions of the environment maps - i.e. the sky above for environment 1 (and also necessarily environment 3). The main source of within-environment variance that makes the odd-one-out task in this study challenging is the variation introduced from changing camera angle. Thus, the success of models looking at the lower part of the object implies that the lower region of the lighting environments is more ‘rotationally’ symmetric, giving more reliable information to perform the odd-one-out task. To validate this idea, we computed the variation of chromatic statistics across camera angles for the upper part and lower part of the object as summarized in Figure 7. To get these variations, for each condition (e.g. glossy and environment 1), we first computed a lightness-weighted mean chromaticity of a gray object (distractor stimuli) for each of 6 camera angles in *L*\**a*\**b*\* space (white circles in right inserted plot) and computed the average across 6 values (red dot). Then, we calculated the average deviation between this averaged color and the lightness-weighted mean of each camera angle (average length across 6 black six lines between white circles and red dot), which gives a measure of the variation of lightness-weighted mean across camera angles. As seen, the upper 75% of the object exhibits larger variation across camera angles than the lower 25% region for both matte and glossy conditions in environment 1 and 3. In contrast, in environment 2, variation is nearly the same for upper and lower regions of the object and also for matte and gloss conditions. These are the variation of lightness-weighted mean chromaticity, but we confirmed similar trends for the variation of mean chromaticity. To sum, these analyses support the idea that the lower region of the lighting environments 1 and 3 is more symmetric across camera angles than the upper region. It is important to note that this trend depends on the content of the specific lighting environment, and to conclude how generally this observation holds in the real world would require systematic analysis of other types of lighting environments (e.g. indoor environment or different weather conditions). Furthermore, this analysis confirmed that when the lower 25% of the object exhibits smaller chromatic variation for glossy objects, it does so also for matte objects, which might be a potential reason why occlusion sensitivity maps were similar between matte and glossy objects.

**Fig. 7.**
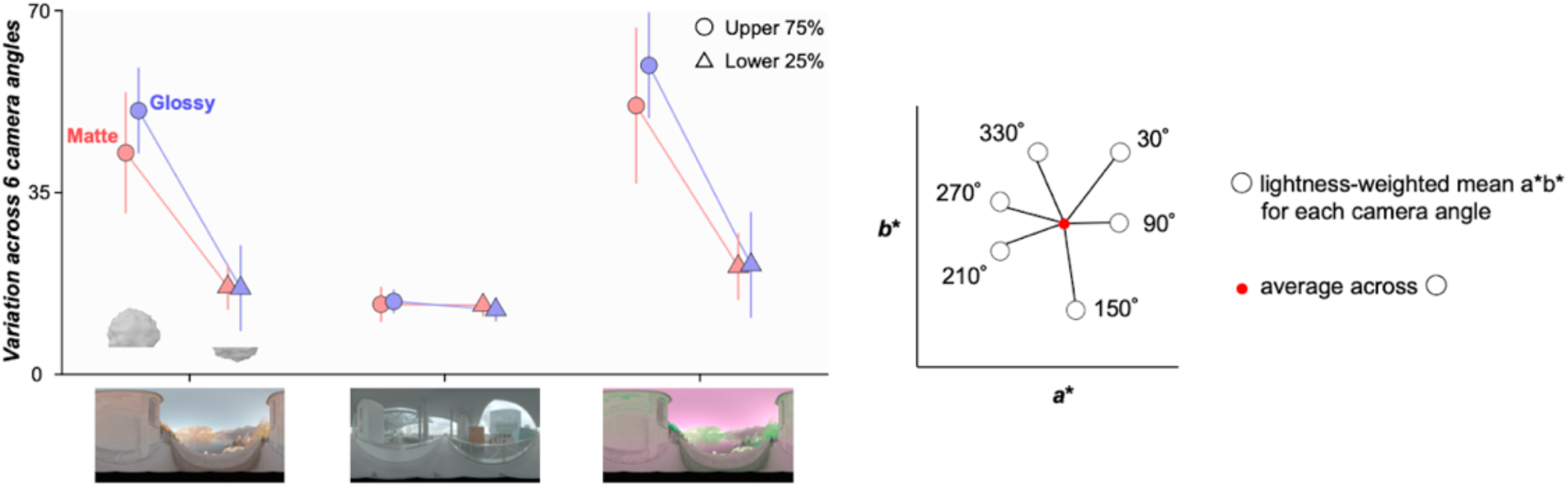
Variation of lightness-weighted mean chromaticity over 6 camera angles for the upper 75% region of the object and for the lower 25 % region of the object. For each condition (e.g. glossy, environment 1 and upper 75%), we computed a lightness-weighted mean color across 6 camera angles, and then calculated the average color difference between this mean color and the mean at each camera angle.

### 3.4 Evaluation of CNNs in novel lighting environments

One limitation in the CNN implementation so far is that CNNs were trained under only three lighting environments, and in the model evaluation stage, chromatic thresholds were estimated under the environments that the CNNs saw during the training process, even though we did use novel, unseen object images presented from different camera angles. This was a necessary choice because the major purpose in this study was to train CNNs using images labeled by human responses and the number of human responses was limited. Nevertheless, it would be desirable to test how well CNNs generalize to novel lighting environments. To test this idea, we first trained neural networks under two of the three environments, and tested under the third environment. However, under such training, CNNs trained by ground-truth and by human responses showed poor performances, and chromatic thresholds could not be estimated in most conditions. Secondly, we further rendered new datasets under 15 novel lighting environments ([28, 13]; and https://hdrmaps.com/freebies/) that cover a diverse range of natural lighting environments. Then, we evaluated the CNNs trained by human responses and ground-truth (ResNet50 presented in the Results section) using the procedure detailed in subsection 2.3. For four environments (labelled A-D) for which it was possible to estimate thresholds, the areas of the resultant discrimination ellipses are summarized in Figure 8. For environments A and B, the human-trained CNN showed better performance (smaller area), showing better generalization to novel environments for both matte and glossy conditions. For the environment C, however, the ground-truth trained CNN showed better performance. For the environment D, the human-trained CNN performed better than the ground-truth CNN for the matte condition, but the trend was reversed for the glossy condition. These results suggest that the level of generalization of CNNs to novel lighting environments depends strongly on the lighting environment, and there are differences in the way that human-trained and ground-truth CNNs generalize to novel environments.

**Fig. 8.**
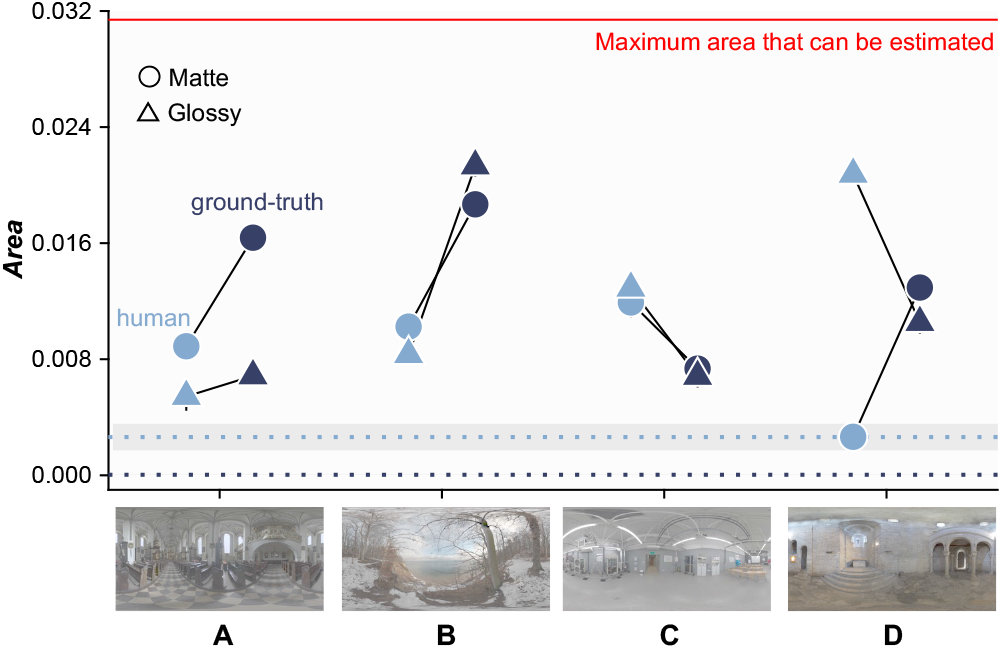
Performance of the human-trained and ground-truth trained CNNs when tested in 15 novel lighting environments. Thresholds could be estimated in 4 environments (A-D), for which the areas of the discrimination ellipses are plotted; in the remaining 11 environments discrimination thresholds fell beyond the gamut defined by realizable natural reflectance samples (shown by red line). Light blue data points show human-trained CNNs, and dark blue data points show ground-truth CNNs. Error bars show ± 1.0 S.E. across 20 sessions. For comparison, average ellipse areas estimated from ResNet18 trained and tested under the same lighting environments are plotted as dotted lines (light blue for human labels and dark blue for ground-truth labels, with the associated shaded regions showing ± 1.0 S.E. across 6 conditions (2 gloss levels × 3 environments)).

## 4. General Discussion

The major purpose of this study was to model surface color discrimination of matte and glossy objects within each of three lighting environments. Under these conditions, the rendered objects exhibit large color variations over the proximal image of the surface. Two approaches were taken: the first approach used a variety of chromatic statistics extracted from the object region of the proximal image, and another utilized the recent advances in machine learning algorithms to produce trained CNNs to perform the task. Hand-crafted models provided us with several insights. First, none of the tested chromatic statistics models are individually sufficient to achieve human-like discrimination behavior across all conditions, though heuristic effectiveness is condition-dependent. For example, the brightest model generally predicted human thresholds better than the other models in matte conditions, however it performed much worse than the other models and humans in glossy conditions. This was to be expected, as glossy objects feature specular reflections which are chromatically unmodified representations of the environmental illumination. It follows that under this model bright regions of specular reflection, unrepresentative of object color, were selected for comparison, leading to high discrimination thresholds. Both the mean chromaticity and mean chromaticity weighted by lightness or luminance models were shown to be effective in predicting thresholds in environments 1 and 3. The machine learning approach revealed that CNNs trained by human responses can reproduce human thresholds nearly perfectly whilst CNNs trained by ground-truth labels showed much better performance than humans, leading to high RMSE values. We further used a technique to visualize the region of an object that drives a network to perform the discrimination task and identified that the lower region of the object is key to performing surface color discrimination for both matte and glossy objects. When chromatic statistics models were also restricted to this region of the objects, they more closely predicted human performance.

One major bottleneck in hypothesis-based modeling approaches is that researchers have to predict in advance what information could be used by humans. In contrast, behaviors observed in some visual experiments could be complex, as in the case of this study, making it hard to predefine potential strategies. In this regard, a hypothesis-free data-driven approach based on CNNs provides a powerful alternative as CNNs automatically learn to use information useful in reproducing human responses. Because of its computational power, the performance can match human thresholds with minimum error, and reverse engineering of such CNNs could provide insight into underlying computations. CNNs have been used presumably in almost all domains for computer vision by now, but in these engineering applications networks are exclusively trained on ground-truth labels. Recent years have seen increasing use of CNNs in perceptual studies, such as gloss perception [29-31], translucency perception [32], and physical properties of cloth [33]. To our knowledge, however, this is the first time human color discrimination has been modeled based on CNNs trained on human behavioral patterns. Identifying the surface color of an object requires perceptual separation of contributions from lighting and surface properties, but this is a hard ill-posed problem. Our approaches revealed potential strategies humans might take to separate contributions of lighting and materials from the proximal image of an object in a simplistic way that utilizes the regularities of natural environments. Occlusion sensitivity analysis revealed a stronger influence of the lower part of the object’s surface on classification behavior of the CNN. This might be beneficial for judging glossy objects since specular reflection tends to locate around the upper part of the object’s surface. In addition, such strategy was shown to be effective to perform an odd-one-out task specifically in this study because chromatic variation across camera angles for the lower part of objects is lower than that of the upper part of objects in environments 1 and 3. Past studies have used simplistic stimuli in color discrimination. While these stimuli helped researchers to precisely characterize fundamental abilities to discriminate two homogeneously colored matte surfaces, our findings strongly suggest that the discrimination of real-world objects requires additional strategies to disentangle the contributions of light and material.

One caveat of this study is to acknowledge the difficulty in interpreting internal computations in CNNs. Several researchers have discussed the validity of network architecture regression to human cytoarchitecture, such as Hasson et al. [34], who draw a comparison between macroevolution across generations of animals and backpropagation across trials for machine learning algorithms; with functional adaptations in the former being analogous to optimization in the latter. Kriegseskorte [35] has pointed out the emergence of simple orientation-based filters in early layers of CNNs, and more complex object-based filters in later layers of networks trained on visual discrimination and classification tasks that are analogous to simple cells early in the visual system and the inferior temporal lobe in the later parts of the ventral stream. This correspondence is particularly pronounced when stimuli and networks are sufficiently complex, as is the case in our task. These proposals seem to suggest that network architecture, when appropriately trained, organizes itself like neural structures, and therefore can be used as an analogy to the brain when investigating human behavior. Saxe et al. [36] has been skeptical of this, noting the way in which data-driven models can serve to shift the focus of study away from developing specific theories of neural function. Machine learning, they argue, is just one approach in a broader toolkit that psychologists and neuroscientists can use to examine neural functions and should be combined with hypothesis-based testing in humans. Accordingly, while the results of the present study are insufficient to make resolute conclusions about what information humans extract from the proximal image of an object, they provide a foundation for future hypothesis-based testing in humans. The exploration of other techniques to formalize the strategies of CNNs is rapidly expanding, which allows for a further analysis of regions of interest at different layers throughout a network. Future studies using CNNs can benefit from these innovations and may inspire novel hypothesis-based testing.

Overfitting of CNNs to the properties of the training dataset is a well known issue. In this study, we used unseen images, from novel camera angles, to test the model performance. However, given the limited availability of human psychophysical responses for training, we were limited to a small number of lighting environments, and it is possible that features of the models were driven by specific features of these environments. The fact that the human-trained and ground-truth trained networks differed in their patterns of generalization, again suggests that they encoded different regularities that drove their responses. Tests of generalization provide a further method by which to uncover the statistical regularities in the training set that are used by human and machine-learning algorithms.

This study prompts several future research questions. For example, removing regions of interest in a psychophysical study would allow us to confirm a correspondence between the regions significant to the network and to humans during color discrimination. In such an ‘Image Perturbation’ paradigm [37, 38], noise is added to regions of interest according to occlusion sensitivity and thresholds are measured in both pre-trained networks and human participants. Observing meaningful deficits in performance, and concordance between human and network performance after perturbation would help confirm that the networks are performing the task using information that is also important to human observers.

To summarize, the present study demonstrated that image chromatic statistics, especially computed after image segmentation, could be useful in predicting human color discrimination, although their effectiveness is dependent on the types of surfaces being observed, and the types of environments in which they are seen. This study also showed the effectiveness of deep learning techniques in modeling color vision functions, especially when trained using human responses. Networks trained on ground-truth labels perform better than humans, and this makes them less useful for modeling human behavior. Analysis of the most successful networks identifies image regions that are key to reproducing human discrimination thresholds, which prompts future study to empirically test this dependence via psychophysics.

## Acknowledgments

TM is supported by a Sir Henry Wellcome Postdoctoral Fellowship from Wellcome Trust (218657/Z/19/Z) and a Junior Research Fellowship from Pembroke College, University of Oxford. SP is supported by an Oxford-MRC Doctoral Training Partnership, Hoare Lea, LLC and Pembroke College, Oxford. For the purpose of open access, the author has applied a CC BY public copyright license to any Author Accepted Manuscript version arising from this submission.

## Supplementary material

Figure S1 shows RMSE for all tested models in this study. Generally speaking, image segmentation models shown by light green symbols perform well in environments 1 and 3 while for environment 2 1-D chromatic statistics models are the best predictors. These results emphasize that different strategies are required to model human behaviors depending on whether specular reflections are present or not and on the lighting environment in which objects are embedded.

**Figure S1:**
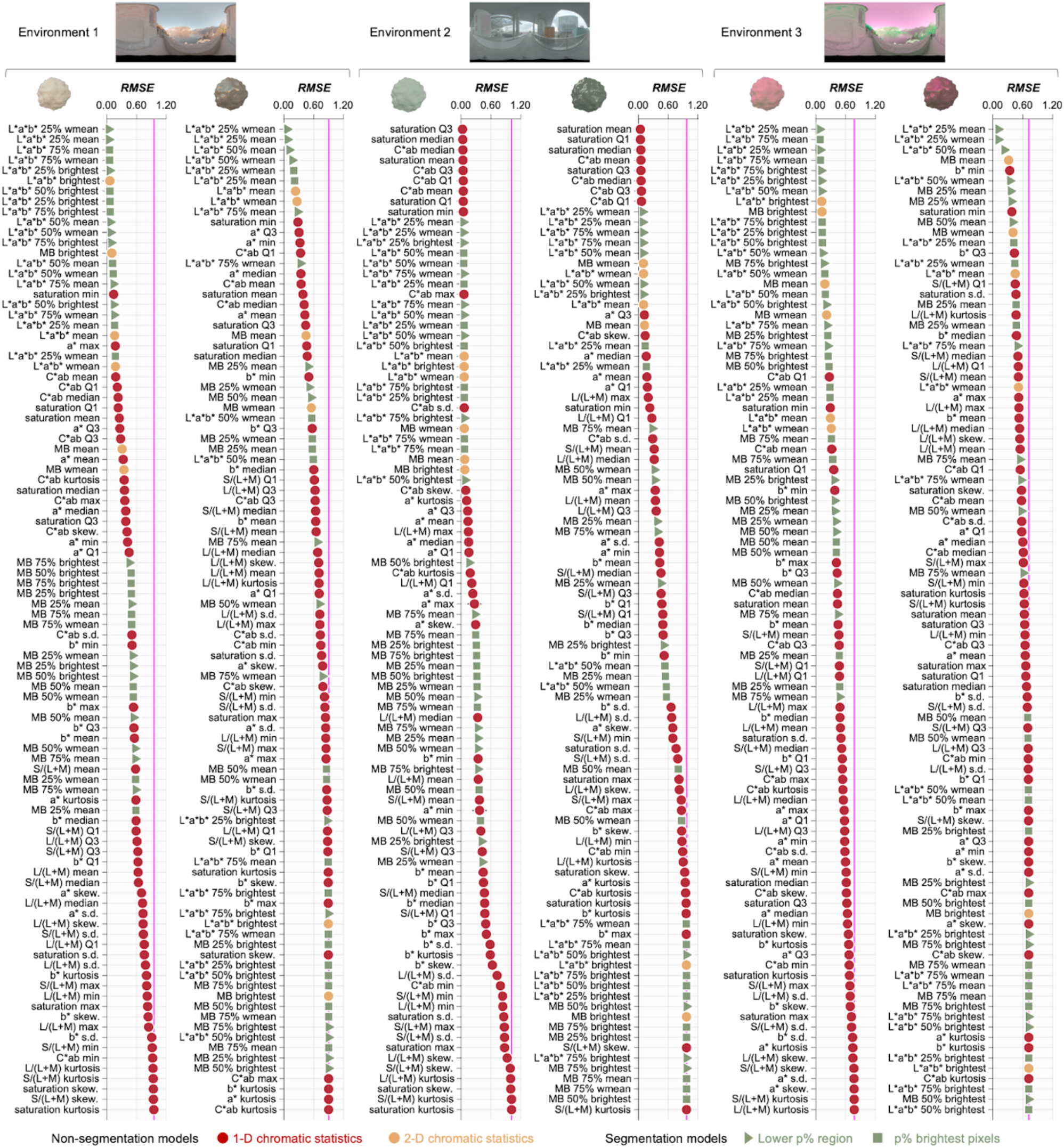
Performance of all chromatic statistics models tested in this study sorted by RMSE values in predicting human performance. Vertical magenta lines show models whose chromatic thresholds could not be estimated for any hue direction because discrimination thresholds fell beyond the gamut defined by realizable natural reflectance samples.

